# Genetic stratification of depression in UK Biobank suggests a subgroup linked to age of natural menopause

**DOI:** 10.1101/134601

**Authors:** David M. Howard, Lasse Folkersen, Jonathan R. I. Coleman, Mark J. Adams, Kylie Glanville, Thomas Werge, Saskia P. Hagenaars, Buhm Han, David Porteous, Archie Campbell, Toni-Kim Clarke, Gerome Breen, Patrick F. Sullivan, Naomi R. Wray, Cathryn M. Lewis, Andrew M. McIntosh

**Affiliations:** Social, Genetic and Developmental Psychiatry Centre, Institute of Psychiatry, Psychology & Neuroscience, King’s College London, UK; Division of Psychiatry, University of Edinburgh, Royal Edinburgh Hospital, Edinburgh, UK; Lundbeck Foundation Initiative for Integrative Psychiatric Research, iPSYCH, Copenhagen, Denmark; Institute of Biological Psychiatry, Mental Health Services Capital Region of Denmark, Copenhagen, Denmark; NIHR Maudsley Biomedical Research Centre, South London and Maudsley NHS Trust, London, UK; Department of Clinical Sciences, University of Copenhagen, Copenhagen, Denmark; Lundbeck Foundation’s Center for GeoGenetics, GLOBE Institute, University of Copenhagen, Denmark; Department of Biomedical Sciences, Seoul National University College of Medicine, Seoul, Republic of Korea; Centre for Genomic and Experimental Medicine, University of Edinburgh, Edinburgh, UK; Centre for Cognitive Ageing and Cognitive Epidemiology, University of Edinburgh, Edinburgh, UK; Usher Institute for Population Health Sciences and Informatics, University of Edinburgh, Edinburgh, UK; Department of Medical Epidemiology and Biostatistics, Karolinska Institutet, Stockholm, Sweden; Department of Genetics, University of North Carolina, Chapel Hill, NC, USA; Department of Psychiatry, University of North Carolina, Chapel Hill, NC, USA; Queensland Brain Institute, University of Queensland, Brisbane, Queensland, Australia; Department of Psychology, University of Edinburgh, Edinburgh, UK

## Abstract

Depression is a common and clinically heterogeneous mental health disorder that is frequently comorbid with other diseases and conditions. Stratification of depression may align sub-diagnoses more closely with their underling aetiology and provide more tractable targets for research and effective treatment. In the current study, we investigated whether genetic data could be used to identify subgroups within people with depression using the UK Biobank. Examination of cross-locus correlations was used to test for evidence of subgroups by examining whether there was clustering of independent genetic variants associated with eleven other complex traits and disorders in people with depression. We found evidence of a subgroup within depression using age of natural menopause variants (*P* = 1.69 × 10^−3^) and this effect remained significant in females (*P* = 1.18 × 10^−3^), but not males (*P* = 0.186). However, no evidence for this subgroup (*P* > 0.05) was found in Generation Scotland, iPSYCH, a UK Biobank replication cohort or the GERA cohort. In the UK Biobank, having depression was also associated with a later age of menopause (beta = 0.34, standard error = 0.06, *P* = 9.92 × 10^−8^). A potential age of natural menopause subgroup within depression and the association between depression and a later age of menopause suggests that they partially share a developmental pathway.

## Introduction

Depression is a common mental health disorder characterised by persistent feelings of sadness or a loss of interest in day-to-day activities lasting for at least a two-week period. These feelings can be accompanied by tiredness, changes in appetite, changes in sleep patterns, reduced concentration, feelings of worthlessness or hopelessness, and thoughts of self-harm or suicide. Zimmerman et al. [1] found that there were 170 different symptom profiles amongst 1566 participants diagnosed with major depressive disorder from the Rhode Island MIDAS project. This variety of different symptom profiles suggest that depression is highly heterogeneous [2]. Depression is also comorbid with many diseases including cancer [3], cardiovascular disease [4] and other psychiatric illnesses [5]. Stratification of depression, to address heterogeneity and comorbidity, may aid in providing valuable aetiological insights and improve treatment efficacy.

Studies aimed at stratifying depression have examined differences between melancholic and atypical depression [6], differences between the sexes and recurrence of the disorder [7] and used data from other traits, such as neuroticism [8] and social contact [9] to stratify depression. Twin-based studies [10] and genome-wide association studies [11, 12] have shown depression to be heritable and genetically correlated with a number of other traits and disorders. This shared genetic component could be due to pleiotropic variants shared across all individuals but could also be as a result of a subgroup for the other trait within depression cases. For example, there is a genetic correlation of −0.11 (standard error = 0.03) between depression and age of natural menopause [13]. If this genetic correlation was due to pleiotropy, then several of the age of menopause variants would be carried by most depression cases. However, if this correlation was due to a subgroup, then a greater proportion of the age of menopause variants would only be carried by individuals in this subgroup. A subgroup could arise where there is a causal association, a shared molecular pathway, a misclassification between the traits, or an ascertainment bias in the diagnosis of depression.

For the current study, BUHMBOX (Breaking Up Heterogeneous Mixture Based On cross(X)-locus correlations) [14] was used to determine whether there was evidence of a subgroup within depression that was genetically more similar to other traits. BUHMBOX uses variants associated with a non-depression trait to calculate weighted pairwise correlations of risk allele dosages within depression cases and controls, adjusted for effect size and allele frequency. Where there is a subgroup amongst depression cases that carry a greater proportion of the risk alleles for the non-depression trait, there will be consistent positive pairwise correlations between those variants (Figure 1). BUHMBOX then calculates a *P*-value based on the likelihood of the observed pairwise correlations between variants.

**Figure 1.**
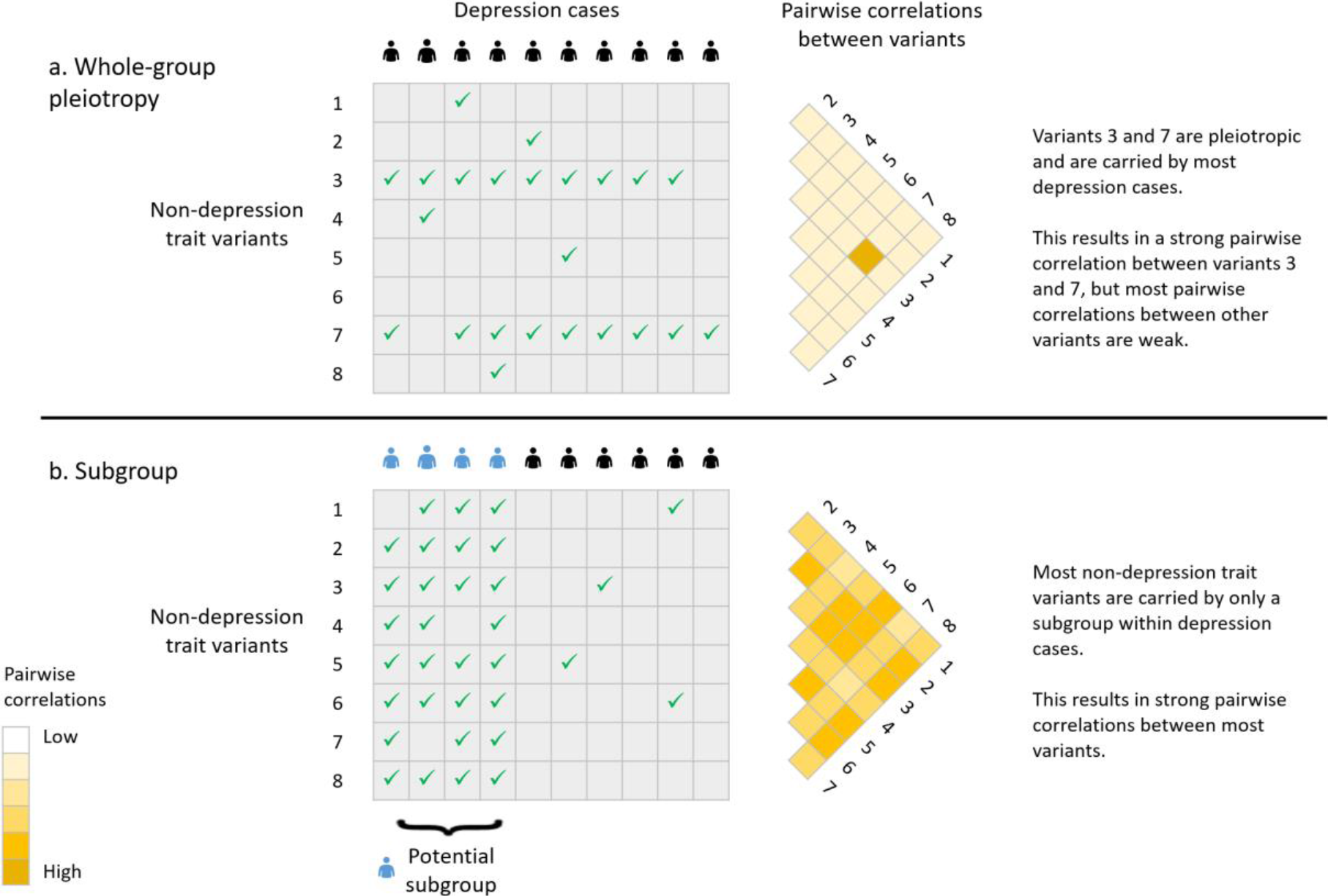
Pairwise correlations between variants for (a) whole-group pleiotropy, where most depression cases carry a few variants associated with a non-depression trait and (b) a subgroup within depression cases 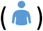, where just the subgroup carry many of the non-depression trait variants. A tick indicates a depression case individual is a carrier of that non-depression variant.

Two definitions of depression were assessed in the UK Biobank [15], one based on the Composite International Diagnostic Interview Short Form (CIDI-SF) [16] and the other based on a broader help-seeking definition (broad depression) [12]. Since many traits are genetically correlated with depression [13], a power calculation was performed to determine traits with sufficient power to detect a subgroup. Power is determined by the number of depression cases, the size of any subgroup within depression cases, the number of associated variants tested from the non-depression trait and the effect sizes of these variants. We tested adequately-powered traits for evidence of a subgroup in depression cases using BUHMBOX v0.38 [14]. Replication of traits forming a subgroup in depression were sought in Generation Scotland: Scottish Family Health Study (GS:SFHS), The Lundbeck Foundation Initiative for Integrative Psychiatric Research (iPSYCH), a UK Biobank replication cohort, and the Genetic Epidemiology Research on Adult Health and Aging (GERA) cohort. UK Biobank and GS:SFHS were used to investigate phenotypic associations between depression and traits forming a subgroup.

## Materials and Methods

### UK Biobank discovery cohort

The UK Biobank is a population-based cohort of 501,726 individuals with imputed genome-wide data for 93,095,623 autosomal genetic variants [15]. A genetically homogeneous sample of 462,065 individuals was identified using the first two principal components from a 4-means clustering approach. A total of 131,790 individuals were identified as being related up to the third degree (kinship coefficients > 0.044) using the KING toolset [17] and were removed from the sample. For these related individuals a genomic relationship matrix was calculated to enable the identification of one individual from each related group that could be reinstated. This allowed the reintroduction of 55,745 individuals providing an unrelated sample of 386,020 individuals.

### UK Biobank depression phenotypes

Two depression phenotypes were assessed for evidence of subgroups in UK Biobank. For the UK Biobank discovery cohort, both phenotypes were restricted to only those individuals that had completed the online mental health questionnaire (n = 109,049). The first phenotype analysed was based on the Composite International Diagnostic Interview Short Form (CIDI-SF) [18] as used by Davis et al. [16] to provide a lifetime instance measure of depression in the UK Biobank. Davis et al. [16] provide a more in-depth description of this CIDI-SF phenotype, but in summary cases were defined as having:

- at least one core symptom of depression (persistent sadness (Data-Field: 20446) or a loss of interest (Data-Field: 20441)) for most or all days over a two-week period which were present “most of the day” or “all of the day”.
- plus at least another four non-core depressive symptoms with some or a lot of impairment experienced during the worst two-week period of depression or low mood.

The non-core depressive symptoms that were included in this assessment of the worst episode of depression were: Feelings of tiredness (Data-Field: 20449), Weight change (Data-Field: 20536), Did your sleep change? (Data-Field: 20532), Difficulty concentrating (Data-Field: 20435), Feelings of worthlessness (Data-Field: 20450), and Thoughts of death (Data-Field: 20437). Cases that self-reported another mood disorder were excluded. Controls were determined by not having at least one core symptom of depression or not endorsing at least another four non-core depressive symptoms if at least one core symptom was endorsed. This provided a total of 25,721 CIDI-SF cases and 61,894 controls.

A second depression phenotype within the UK Biobank discovery cohort was also examined using the broad depression definition from Howard et al. [12] with detailed information provided in that paper. In summary, cases had sought help for nerves, anxiety, tension or depression from either a general practitioner or a psychiatrist (Data-Field: 2090 and Data-Field: 2100), whereas controls had not. Cases were supplemented with an additional 132 individuals identified as having a primary or secondary International Classification of Diseases (ICD)-10 diagnosis of a depressive mood disorder from linked hospital admission records (Data-Field: 41202 and Data-Field: 41204). Participants identified with bipolar disorder, schizophrenia or personality disorder and those reporting a prescription for an antipsychotic medication were removed. This provided a total of 36,790 broad depression cases and 70,304 controls. The phenotypic correlation between the CIDI-SF depression phenotype and the broad depression phenotype was 0.61 with the number of cases and controls shared across the two definitions shown in Supplementary Table 1.

### Traits examined as subgroups within depression

We selected traits genetically correlated with depression (false discovery rate corrected, *q* < 0.01) in Howard et al. [13] to test as subgroups within depression, which included anthropomorphic, autoimmune, life course, cardiovascular and other psychiatric traits. For each trait, there was a requirement that publicly available summary statistics were available and that the UK Biobank was not included in that study due to potential confounding effects (Supplementary Table 2).

The BUHMBOX power calculation test v0.1 [14] was used to determine whether there was sufficient power to detect a subgroup for each depression correlated trait and to identify the optimum variant selection criterion (*P* < 5 × 10^−8^, *P* < 10^−6^ or *P* < 10^−4^). The power calculation was conducted using the CIDI-SF depression phenotype and then using the broad depression phenotype. Variants from the summary statistics for each non-depression trait were examined in the UK Biobank discovery cohort. Variants that had a call rate less than 0.99, were out of Hardy-Weinberg equilibrium (*P* < 10^−10^), had a hard call threshold less than 0.25, or had a minor allele frequency less than 0.005 were excluded. BUHMBOX requires that all variants are available for all individuals and therefore individuals with a call rate less than 1 were removed. To identify independently segregating variants, clumping was conducted in PLINK v1.90b4 [19] using an r^2^ value of 0.01 across a 3Mb window in either CIDI-SF or broad depression control individuals, respectively.

For the power analysis the approach used in Han et al. [14] was followed, with 1000 simulated iterations run for each trait, the proportion of individuals in the subgroup was set to 0.2 and a nominal subgroup *P*-value of 0.05 was used. Power analyses were used to identify the optimum variant selection criterion that provided the greatest power for each non-depression trait. Where power was the same across variant selection criteria, the strictest variant selection criterion was selected as the optimum. Variants with *P* < 10^−4^ were not publicly available for Squamous Cell Lung Cancer or Lung Cancer and so *P* < 10^−5^ was used instead. Only those traits that had a power > 0.8 (using the optimum variant selection criterion) were selected to be tested for evidence of a subgroup within depression.

### Testing for subgroups within depression

For the traits that had power > 0.8, variants meeting the optimum variant selection criterion were extracted from the UK Biobank discovery cohort. The same quality control thresholds and method to identify independently segregating variants as used as previously in the power analysis were applied. BUHMBOX v0.38 [14] was used to examine shared risk alleles for each non-depression trait within CIDI-SF depression and broad depression. BUHMBOX uses the positive correlations between risk allele dosages in cases to determine whether any sharing of risk alleles is driven by all individuals (whole-group pleiotropy) or by a subset of individuals (Figure 1). The likelihood of observing such positive correlations are used to determine the subgroup *P*-values. The BUHMBOX software and manual are freely downloadable from http://software.broadinstitute.org/mpg/buhmbox/.

Sex, age, genotyping array and the first 20 principal components were fitted as covariates in the subgroup analysis. Bonferroni correction was used to account for the multiple testing of non-depression traits, with *P*-values < 5 × 10^−3^ (0.05/10) or < 4.5 × 10^−3^ (0.05/11) deemed significant for CIDI-SF or broad depression, respectively. No multiple testing correction was applied for the two depression definitions analysed.

BUHMBOX calculates and outputs polygenic risk scores for each individual based on the summary statistics provided. If a subgroup for a trait exists, as in Figure 1b, then potentially this subgroup would carry a greater number of these variants compared to the non-subgroup depression cases and therefore a binomial distribution would exist within the polygenic risk scores of cases. To examine whether the standardised distributions of polygenic risk scores for non-depression traits in depression cases and controls could be explained by two univariate normal distributions the mix2normal function from the VGAM package [20] in R v3.5.2 was used. The use of polygenic risk scores provides additional supporting evidence of a subgroup and provides an estimation of the size of any subgroup.

### Replication of significant subgroups within depression

Traits that showed significant evidence of forming a subgroup for depression in the UK Biobank discovery cohort were re-examined in independent cohorts: Generation Scotland: Scottish Family Health Study (GS:SFHS), The Lundbeck Foundation Initiative for Integrative Psychiatric Research (iPSYCH), a UK Biobank replication cohort, and the Genetic Epidemiology Research on Adult Health and Aging (GERA) cohort. In each of the replication cohorts, individuals were removed if they had a variant call rate less than 1 and variants were removed if they had a call rate less than 0.99, were out of Hardy-Weinberg equilibrium (P < 10^−10^) or had a minor allele frequency less than 0.005.

The family and population-based GS:SFHS cohort [21] consisted of 23,960 individuals, of whom 20,195 were genotyped and subsequently imputed [22] providing a total of 8,633,288 variants for 20,032 individuals (11,085 females and 8,947 males). An unrelated subset was created using GCTA v1.22 [23] ensuring that no two individuals shared a genomic relatedness of ≥ 0.025. Individuals were removed if they were identified as population outliers [24] or had participated in UK Biobank (using a checksum-based approach [25]). Sex, age and the first 20 principal components were fitted as covariates in the subgroup tests. A diagnosis of major depressive disorder (MDD) was made using two initial screening questions and the Structured Clinical Interview for the Diagnostic and Statistical Manual of Mental Disorders [26] and has been described previously in Fernandez-Pujals et al. [27]. Using record linkage to the Scottish Morbidity Record, we removed 1,072 controls who had attended at least one psychiatry outpatient clinic. Using the psychiatric inpatient records, we identified 47 MDD cases who were also diagnosed with bipolar disorder or schizophrenia and these individuals were excluded. This provided a total of 975 MDD cases and 5,971 controls. These participants provided prior consent for their anonymised data to be linked to medical records.

iPSYCH is a case-control sample with genotyping data collected for 77,639 individuals after quality control. The iPSYCH sample was phased using SHAPEIT3 [28] and imputed using Impute2 [29] using the 1000 genomes phase 3 data [30]. An unrelated subsample was identified using the KING toolset
[17] with second degree relatives or closer excluded. Sex, age, genotyping array and the first 20 principal components were fitted as covariates in the subgroup tests. Depression status was ascertained from in- and out-patient hospital records with controls screened to ensure they had no other psychiatric disorders. This provided a total of 19,644 cases and 21,295 controls. Further detailed information on the iPSYCH sample is available in Pedersen et al. [31].

The UK Biobank discovery sample consisted of only those individuals that completed the mental health questionnaire. Therefore, the individuals that did not complete the questionnaire were used as an independent replication cohort. For these individuals only the broad depression definition could be assessed and applying the same quality control criteria used in the UK Biobank discovery cohort resulted in 71,282 broad depression cases and 128,303 controls.

The GERA cohort is a genotyped subsample of 78,419 participants from the Kaiser Permanente Medical Care Plan, Northern California Region (https://www.ncbi.nlm.nih.gov/projects/gap/cgi-bin/study.cgi?study_id=phs000674.v3.p3). GERA was genotyped using custom designed Affymetrix Axiom arrays [32] before being phased with SHAPEIT v2.5 [33] and imputed with IMPUTE2 v2.3.1 [29] using the 1000 Genomes Project [30] as the reference panel. GERA is a mixed ancestry cohort [34] and in the current analysis only those individuals with European ancestry were examined with related individuals up to the third degree removed. Sex, age and the first 10 principal components were fitted as covariates in the subgroup tests. MDD status was ascertained using ICD-9 coding from in- and out-patient hospital records which identified 4,912 MDD cases and 33,902 controls.

During the replication analysis, the BUHMBOX power calculation test v0.1 [14] was applied assuming the same estimated proportion of individuals in the subgroup and the same variant selection criterion as in the UK Biobank discovery cohort, and a nominal subgroup *P*-value of 0.05. BUHMBOX v0.38 [14] was then used to examine whether there was evidence of a subgroup within depression cases in the GS:SFHS, iPSYCH, UK Biobank replication and GERA cohorts. The power and subgroup analyses were run using all individuals and then, as age of natural menopause trait generated a significant result in the subgroup analysis the analyses, were run using females only.

### Phenotypic examination of significant subgroups within depression

The age of natural menopause trait generated a significant result in the subgroup analysis in the UK Biobank discovery cohort and we therefore examined whether those with depression had a later or earlier onset of menopause compared to controls. This was conducted in UK Biobank and GS:SFHS using unrelated individuals by applying the same criteria as described previously to identify relatedness. In UK Biobank, a linear regression was conducted to compare the age of menopause (Data-Field: 3581) in the CIDI-SF depression cases with controls covarying for the age when attending assessment centre (Data-Field: 21003). Age of attending assessment centre was fitted as a covariate as it was associated with age of menopause (beta = 0.12, standard error = 4.92 × 10^−3^, *P* = 10^−137^). Individuals that reported they had not experienced menopause, were unsure whether menopause had been experienced or had undergone a hysterectomy were excluded. The latest entry for each individual, at either the Initial Assessment (2006-2010), Repeat Assessment (2012-2013) or Imaging visit (2014+), was used to record the age of menopause and age when attending assessment centre. Individuals that had an age at onset of depression (Data-Field: 20433) that was two years prior to or after the age of menopause were classified as controls. In GS:SFHS, a linear regression was also used to compare the age of menopause in MDD cases and controls covarying for age when attending assessment centre. Individuals that had a self-reported age at onset of first episode of MDD [27] obtained during the Structured Clinical Interview that was two years prior to or after the age of menopause were classified as controls. GS:SFHS individuals that reported they had not experienced menopause, had a hysterectomy or whose ovaries had been removed were excluded.

## Results

### Power analyses of potential subgroups traits

To determine whether there was sufficient power (> 0.8) to detect a subgroup and identify the optimum variant selection criterion (*P* < 5 × 10^−8^, *P* < 10^−6^ or *P* < 10^−4^) for each trait the BUHMBOX power calculation test v0.1 [14] was used. The results of the power analysis for detecting a subgroup for 25 available traits within the two depression definitions are provided in Table 1. Obesity 1 and Obesity 3 were from the same study [35] and were highly correlated (r_g_ = 0.942, standard error = 0.045) and therefore only the trait providing greatest power (Obesity 3) was selected to be tested as a subgroup. The same approach was used for the Squamous Cell Lung Cancer and Lung Cancer traits with only Squamous Cell Lung Cancer selected for analysis.

**Table 1.** Power analysis for detecting a subgroup for 25 traits within either Composite International Diagnostic Interview Short Form (CIDI-SF) depression or broad depression in the UK Biobank discovery cohort. PubMed identifiers (PubMed ID) for the 25 traits are provided. Bold values indicate that power was > 0.8. The optimum variant selection criterion that maximised power for the non-depression traits are provided. †Variants with *P* < 10^−4^ were not publicly available for Squamous Cell Lung Cancer or Lung Cancer and so *P* < 10^−5^ was tested instead.

| Subgroup trait | PubMed ID | CIDI-SF depression |  | Broad depression |  |
| --- | --- | --- | --- | --- | --- |
|  |  | Optimum variant selection criterion | Power | Optimum variant selection criterion | Power |
| Neuroticism | 24828478 | $< 10^{-4}$ | 0.197 | $< 10^{-4}$ | 0.212 |
| Schizophrenia | 25056061 | $< 10^{-4}$ | **1** | $< 10^{-6}$ | **1** |
| Bipolar Disorder | 29906448 | $< 10^{-4}$ | **1** | $< 10^{-4}$ | **1** |
| Attention Deficit Hyperactivity Disorder | 27663945 | $< 10^{-4}$ | 0.319 | $< 10^{-4}$ | 0.416 |
| Autism Spectrum Disorder | 28540026 | $< 10^{-4}$ | **1** | $< 10^{-4}$ | **1** |
| Anorexia Nervosa | 28494655 | $< 10^{-4}$ | **1** | $< 10^{-4}$ | **1** |
| Triglyceride Level | 24097068 | $< 10^{-4}$ | 0.265 | $< 10^{-4}$ | 0.364 |
| Coronary Artery Disease | 26343387 | $< 10^{-4}$ | **0.986** | $< 10^{-4}$ | **1** |
| Crohn's Disease | 26192919 | $< 5 \times 10^{-8}$ | **1** | $< 5 \times 10^{-8}$ | **1** |
| Inflammatory Bowel Disease | 28067908 | $< 5 \times 10^{-8}$ | **1** | $< 5 \times 10^{-8}$ | **1** |
| Waist to Hip Ratio | 25673412 | $< 10^{-4}$ | 0.114 | $< 10^{-4}$ | 0.117 |
| Body Fat | 26833246 | $< 10^{-4}$ | 0.257 | $< 10^{-4}$ | 0.325 |
| Waist Circumference | 25673412 | $< 10^{-4}$ | 0.143 | $< 10^{-4}$ | 0.143 |
| Overweight | 23563607 | $< 10^{-4}$ | 0.423 | $< 10^{-4}$ | 0.551 |
| Obesity 1 | 23563607 | $< 10^{-4}$ | **0.977** | $< 10^{-4}$ | **0.996** |
| Obesity 3 | 23563607 | $< 10^{-4}$ | **1** | $< 10^{-4}$ | **1** |
| Body Mass Index | 25673413 | $< 10^{-4}$ | 0.135 | $< 10^{-4}$ | 0.14 |
| Age of Menarche | 25231870 | $< 10^{-4}$ | 0.461 | $< 10^{-4}$ | 0.537 |
| Age of Natural Menopause | 26414677 | $< 5 \times 10^{-8}$ | **1** | $< 5 \times 10^{-8}$ | **1** |
| Years of Schooling | 25201988 | $< 10^{-4}$ | 0.081 | $< 10^{-4}$ | 0.108 |
| College Completion | 25201988 | $< 10^{-4}$ | 0.602 | $< 10^{-4}$ | 0.672 |
| Ever Smoked | 20418890 | $< 10^{-4}$ | 0.733 | $< 10^{-4}$ | **0.842** |
| Age of Smoking Initiation | 20418890 | $< 10^{-4}$ | 0.079 | $< 10^{-4}$ | 0.069 |
| Squamous Cell Lung Cancer† | 28604730 | $< 10^{-5}$ | **0.999** | $< 10^{-5}$ | **0.999** |
| Lung Cancer† | 28604730 | $< 10^{-5}$ | **0.813** | $< 10^{-5}$ | **0.889** |

Ten traits had power > 0.8 across both the CIDI-SF depression and broad depression definitions: Schizophrenia [36], Bipolar Disorder [37], Autism Spectrum Disorder [38], Anorexia Nervosa [39], Coronary Artery Disease [40], Crohn’s Disease [41], Inflammatory Bowel Disease [42], Obesity 3 [35], Age of Natural Menopause [43], and Squamous Cell Lung Cancer [44]. There was one further trait, Ever Smoked [45], that had power > 0.8 for detection of a subgroup in broad depression.

### Testing for subgroups within depression

BUHMBOX v0.38 [14] was used to test ten traits for evidence of a subgroup within CIDI-SF depression and eleven traits within broad depression. The results of the subgroup analyses are provided in Table 2. There was evidence of a genetic subgroup relating to age of natural menopause within CIDI-SF depression (*P* = 1.69 × 10^−3^) which remained significant after correction for multiple testing. The 47 variants used to identify this subgroup are provided in Supplementary Table 3. A genetic subgroup relating to age of menopause was detected within the broad depression phenotype (*P* = 9.13 × 10^−3^), although this was not significant after correction for multiple testing.

**Table 2.** Evidence of a subgroup from traits tested within either Composite International Diagnostic Interview Short Form (CIDI-SF) depression or broad depression in the UK Biobank discovery cohort. The number of individuals in the UK Biobank discovery cohort classified as depression cases and depression controls is provided. The number of variants assessed is provided based on the optimum variant selection criterion for that trait. Bold values indicate significant evidence of a subgroup after Bonferroni correction for multiple testing.

| Depression definition | Subgroup trait | Number of variants | Depression cases | Depression controls | Subgroup P-value |
| --- | --- | --- | --- | --- | --- |
| CIDI-SF | Schizophrenia | 1,107 | 1,053 | 2,393 | 0.48 |
|  | Bipolar Disorder | 441 | 8,027 | 19,146 | 0.54 |
|  | Autism Spectrum Disorder | 56 | 20,524 | 49,050 | 0.94 |
|  | Anorexia Nervosa | 180 | 15,233 | 36,519 | 0.96 |
|  | Coronary Artery Disease | 305 | 12,941 | 31,186 | 0.28 |
|  | Crohn's Disease | 62 | 23,233 | 55,940 | 0.31 |
|  | Inflammatory Bowel Disease | 146 | 19,534 | 46,942 | 0.22 |
|  | Obesity 3 | 61 | 22,096 | 53,312 | 0.50 |
|  | Age of Natural Menopause | 47 | 23,592 | 56,764 | **$1.69 \times 10^{-3}$** |
|  | Squamous Cell Lung Cancer | 52 | 22,677 | 54,693 | 0.48 |
| Broad | Schizophrenia | 184 | 21,451 | 41,109 | 0.30 |
|  | Bipolar Disorder | 440 | 11,368 | 21,842 | 0.59 |
|  | Autism Spectrum Disorder | 56 | 29,279 | 55,882 | 0.86 |
|  | Anorexia Nervosa | 180 | 21,804 | 41,519 | 0.95 |
|  | Coronary Artery Disease | 305 | 18,514 | 35,428 | 0.31 |
|  | Crohn's Disease | 62 | 33,137 | 63,567 | 0.11 |
|  | Inflammatory Bowel Disease | 146 | 27,948 | 53,235 | 0.12 |
|  | Obesity 3 | 61 | 31,690 | 60,548 | 0.35 |
| | Age of Natural Menopause | 47 | 33,685 | 64,533 | $9.13 \times 10^{-3}$ |
|  | Squamous Cell Lung Cancer | 52 | 32,535 | 62,071 | 0.85 |
|  | Ever Smoked | 99 | 27,521 | 52,296 | 0.34 |

Density plots of the distributions of standardised polygenic risk scores, calculated using 47 variants with *P* < 10^−4^ for age of natural menopause, in CIDI-SF depression cases and controls with density curves of the estimates for underlying univariate normal distributions are provided in Figure 2. In CIDI-SF depression cases, one normal distribution had a mean polygenic risk score of −0.11 (standard deviation = 0.75) with a second normal distribution with a mean of 0.86 (standard deviation = 0.76). The proportion of individuals in the second normal distribution was 0.11 which is potentially indicative of the proportion of case individuals in the age of menopause subgroup. Cohen’s d was greater for the two univariate distributions in CIDI-SF depression cases (1.3) than for the controls (0.5).

**Figure 2.**
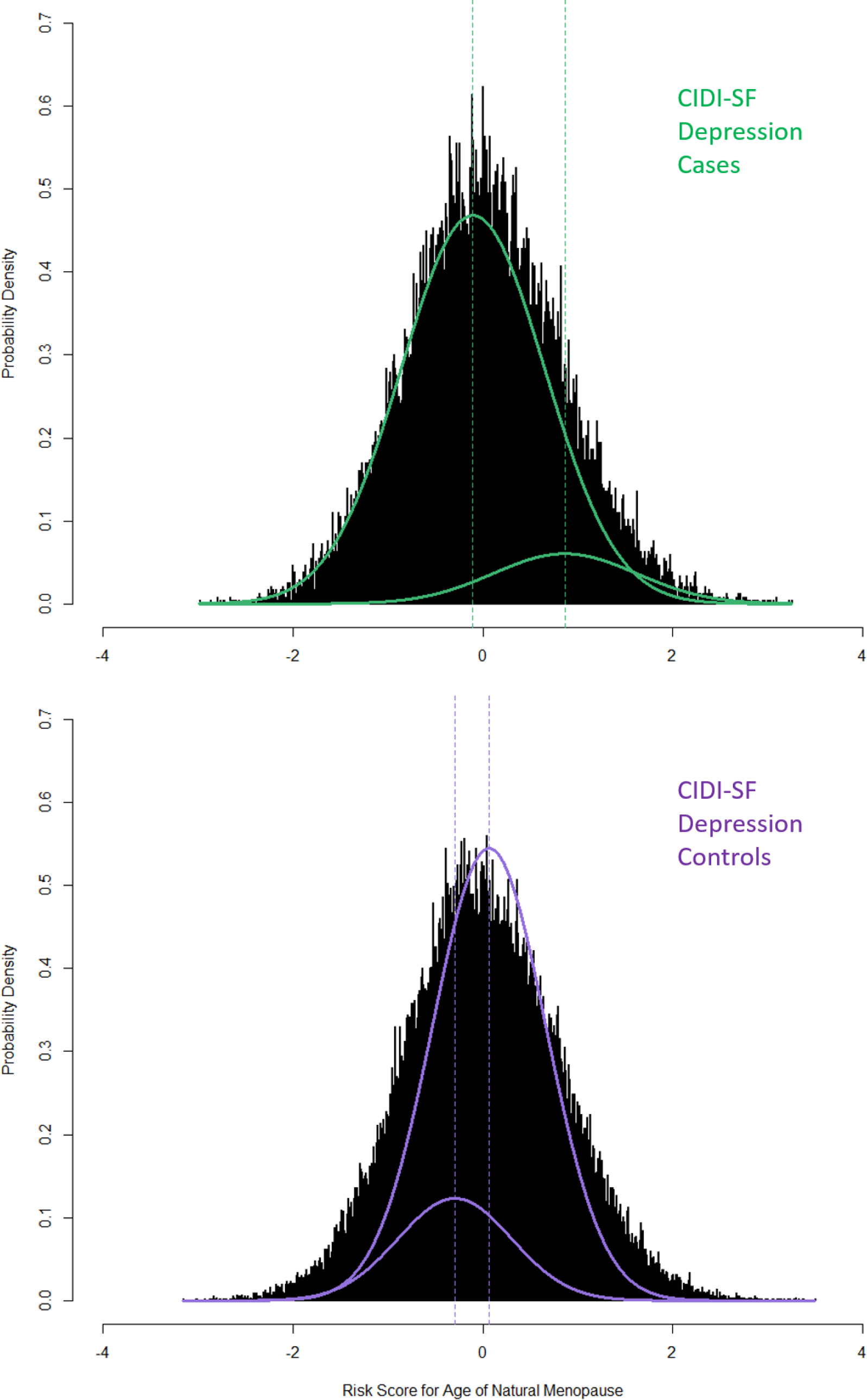
Density plots of the distributions of polygenic risk scores for age of natural menopause in Composite International Diagnostic Interview Short Form (CIDI-SF) depression cases and controls. Overlaid density curves are used to provide estimates of underlying univariate normal distributions in cases (green) and controls (purple).

As a subgroup was observed for age of natural menopause, which is a sex-limited trait, the subgroup analysis was rerun in men and woman separately, using variants with *P* < 5 × 10^−8^. This would potentially reveal whether it was the genetic variants for age of menopause alone, regardless of sex, which indicated a depression subgroup. In males (7408 cases, 28,558 controls), there was no evidence (*P* = 0.186) for an age of menopause subgroup in CIDI-SF depression. In females (cases = 16,184, controls = 28,206), there remained evidence (*P* = 1.18 × 10^−3^) of an age of menopause subgroup within CIDI-SF depression. Using the mix2normal function to examine age of menopause polygenic risk scores for female depression cases estimated one normal distribution with a mean of −0.18 (standard deviation = 0.73) with a second normal distribution with a mean of 0.35 (standard deviation = 0.86) with the proportion of individuals in the first normal distribution estimated as 0.35.

To replicate the age of menopause subgroup within CIDI-SF depression observed in the UK Biobank discovery cohort, we also examined the GS:SFHS, iPSYCH, UK Biobank replication and GERA cohorts. There was no evidence of a subgroup for age of natural menopause in any of the replication cohorts (*P* ≥ 0.05), when analysing both sexes and in the female only analysis (Table 3). The power was greater in the female only analyses compared to both sexes and this was likely due to the potential subgroup size being larger when analysing only females.

**Table 3.** *P*-values for an age of natural menopause subgroup in depression within the Generation Scotland: Scottish Family Health Study (GS:SFHS), The Lundbeck Foundation Initiative for Integrative Psychiatric Research (iPSYCH), UK Biobank replication and the Genetic Epidemiology Research on Adult Health and Aging (GERA) cohorts. The number of variants is based on an optimum variant selection criterion of *P* < 5 × 10^−8^ for an association with age of natural menopause. Power is based on the estimated proportion of individuals in the age of natural menopause subgroup observed in the UK Biobank discovery cohort (0.11 or 0.35 in female only analysis).

| Cohort | Depression definition | Number of variants | Depression cases | Depression controls | Power | Subgroup *P*-value |
| --- | --- | --- | --- | --- | --- | --- |
| GS:SFHS | MDD | 49 | 975 | 5971 | 0.44 | 0.47 |
|  | MDD (female only) | 49 | 683 | 3250 | 0.87 | 0.74 |
| iPSYCH | Depression | 29 | 19644 | 21295 | 0.60 | 0.24 |
|  | Depression (female only) | 29 | 13299 | 10362 | 1 | 0.33 |
| UK Biobank replication | Broad Depression | 44 | 71282 | 128303 | 1 | 0.17 |
|  | Broad Depression (female only) | 45 | 45136 | 59251 | 1 | 0.37 |
| GERA | MDD | 42 | 4912 | 33902 | 0.92 | 0.47 |
|  | MDD (female only) | 42 | 3543 | 19494 | 1 | 0.63 |

### Phenotypic examination of depression and age of menopause

Having observed evidence for a genetic subgroup for age of natural menopause within CIDI-SF depression, we examined whether age of natural menopause differed between depression cases and controls using a linear model. To examine depression prior to onset of menopause the analysis was restricted to cases that reported depression at least two years prior to onset of menopause with age when attending assessment centre (to assess age of menopause) fitted as a covariate in both UK Biobank and GS:SFHS. In UK Biobank, the age of natural menopause in CIDI-SF depression cases (n = 7312, mean = 50.24 years) was significantly later (beta = 0.34, standard error = 0.06, *P* = 9.92 × 10^−8^) than in controls (n = 21,829, mean = 50.09 years). In GS:SFHS, the age of natural menopause in MDD cases (n = 63, mean = 55.0 years) was earlier than in MDD controls (n = 533, mean 59.0), but after covarying for age of assessment the estimate was in the opposite direction (i.e. depression cases had a later age of menopause) and was not significant (beta = 0.87, standard error = 0.68, *P* = 0.20).

## Discussion

Depression is a heterogeneous mental health disorder and is comorbid with many other diseases and illnesses. Over the last few years, valuable progress has been made in understanding the underlying genetic architecture of depression [11, 13, 46]. Furthermore, stratifying depression using genetic data remains a key goal within the psychiatric genetics community [47] and should lead to improved classification of mental health conditions and more efficacious treatment for patients. Machine learning [48, 49] and polygenic risk score [6, 50] approaches offer possible methods for stratification in mental health. In the current study, we used BUHMBOX [14] to identify whether traits that were genetically correlated with depression were correlated due to a subgroup, i.e. the correlation was driven by a subset of depressed individuals who had a greater genetic loading for the trait. Evidence of a subgroup within depression may provide future opportunities for stratifying the disease.

We examined 25 traits genetically correlated with depression using individuals that had completed the UK Biobank mental health questionnaire. Two definitions of depression were examined to allow a direct comparison between a stricter and a broader definition of depression. We initially conducted a power analysis to determine those correlated traits which could be reasonably tested as genetic subgroups. There were ten traits adequately powered to be tested as subgroups within CIDI-SF depression and eleven traits tested as subgroups within broad depression. A genetic subgroup for age of natural menopause was found within CIDI-SF depression after correction for multiple testing. A genetic subgroup for age of natural menopause was also found within broad depression, but this did not survive multiple testing correction. No evidence for this subgroup was found in GS:SFHS, iPSYCH, a UK Biobank replication cohort or GERA. The lack of replication could be due to Type 1 error, there could be something distinct about the UK Biobank discovery cohort, the different definitions of depression examined, or a combination of factors.

From BUHMBOX, it is not directly possible to determine whether an earlier or later age of menopause led to the observed genetic subgroup. However, the phenotypic analyses conducted suggested people with depression have a later age of menopause and so for the purposes of illustrating possible explanations for this subgroup, a later age of menopause is used. Han et al. [14] suggested that subgroups could arise due to ascertainment bias, misclassification, causal relationships, or molecular subgroups. Ascertainment bias seems unlikely as that would require that a later age of menopause somehow increases the chances of individuals receiving clinical attention and obtaining a diagnosis of depression. Misclassification also seems unlikely as there is no obvious reason why individuals with a later age of menopause would be misdiagnosed as depressed. No evidence for a causal relationship in either direction (P = 0.169 for depression being causing for age of menopause and P = 0.529 for age of menopause being causal for depression) was found by Howard et al. [13] using Mendelian randomization. Molecular subtypes, where there exists a shared developmental pathway between a later age of menopause and depression, represents a potential explanation for our results and identifying this pathway could form the basis of future research.

The relationship between depression and menopause has been well studied, but with inconsistent findings [51, 52]. Studies have reported an increase in depressive symptoms during menopause [53–56], but this may be due to the onset of climacteric symptoms, such as insomnia, heavy sweating, hot flashes, and irritability, rather than menopausal state [57–59]. Whereas, Kaufert et al. [60] reported that there was no effect of onset of menopause on depressive status. A meta-analysis of 14 studies found that an older age at menopause led to a lower risk of depression in later life [61]. Several shared neuroendocrine mechanisms have been proposed between menopause and depression. Failure of the gamma-aminobutyric acid A (GABA_A_) receptor to adapt to fluctuations in ovarian hormones due to the menopause may impact hypothalamic pituitary adrenal (HPA) axis activity [62], with dysregulation of the HPA axis associated with depression [63]. Further, oestradiol is a reproductive hormone that declines during menopause, but it also has a neuroprotective role and contributes to the maintenance of brain homeostasis [64]. A review by Rubinow et al. [65] reported that there was some evidence that oestradiol had an antidepressant effect in perimenopausal women. The role of oestradiol throughout the life course may have produced the results observed in the current study with observable effects on both depression and age of menopause.

The results from the subgroup analysis suggest that there was a shared genetic component underlying both depression and age of menopause. Studies examining genetic factors relating to both menopause and mental health phenotypes have principally been focused on the estrogen receptor alpha (*ESR1*) gene [66], with *ESR1* associated with anxiety [67], premenstrual dysphoric disorder [68], and major depressive disorder [69]. However, variants in or near the *ESR1* gene were not associated with age of menopause [43] and therefore not included in the current analysis. Future research identifying genetic factors underlying shared biological mechanisms between menopause and depression may aid in developing new treatments for related mood disorders.

The limitations of the current study include selection bias, whereby particular individuals are more likely to participate in population-based cohorts or complete additional assessments, such as the online mental health questionnaire. Participants of the UK Biobank are healthier and from less deprived areas than the general population[70] and those that completed the mental health questionnaire had a lower genetic predisposition to severe depression than those who did not [71]. UK Biobank participants that either had a self-reported or a hospital diagnosis of schizophrenia or bipolar disorder were excluded in the current analysis which may limit the potential for identifying subgroups for these disorders. The replication cohorts each used different diagnostic criteria for depression and also examined slightly different sets of genetic variants, nevertheless the set of variants examined were associated with age of menopause. Over half of the traits that are genetically correlated with depression were not included in the subgroup analysis due to a lack of power (≤ 0.8). As increasing large genome-wide association studies become available, a greater number of variants will meet the required selection criteria, allowing additional traits to be tested for evidence of a subgroup within depression.

Depression is both polygenic and heterogeneous and stratification of the disorder may lead to improvements in treatment outcomes. In the current study, we found that depressed individuals in the UK Biobank and GS:SFHS had a later age of menopause. This relationship may have a genetic basis with age of natural menopause found to form a subgroup within UK Biobank CIDI-SF depression cases. Using genetic data to identify individuals in this subgroup may ultimately reveal more efficacious treatments for depression.

## Acknowledgements

This research was conducted using the UK Biobank resource, application number 4844. We are grateful to the UK Biobank and all its voluntary participants. The UK Biobank study was conducted under generic approval from the NHS National Research Ethics Service (approval letter dated June 17, 2011, Ref 11/NW/0382). All participants gave full informed written consent.

Generation Scotland received core support from the Chief Scientist Office of the Scottish Government Health Directorates [CZD/16/6] and the Scottish Funding Council [HR03006]. Genotyping of the GS:SFHS samples was funded by the Wellcome Trust (Wellcome Trust Strategic Award “Stratifying Resilience and Depression Longitudinally” (STRADL) Reference 104036/Z/14/Z) and the Medical Research Council UK and was carried out by the Genetics Core Laboratory at the Wellcome Trust Clinical Research Facility, Edinburgh, Scotland. Ethics approval for the Generation Scotland was given by the NHS Tayside committee on research ethics (reference 15/ES/0040), and all participants provided written informed consent for the use of their data.

The iPSYCH team acknowledges funding from the Lundbeck Foundation (grants R102-A9118 and R155-2014-1724), the Stanley Medical Research Institute, the European Research Council (project 294838), the Novo Nordisk Foundation for supporting the Danish National Biobank resource, and Aarhus and Copenhagen Universities and University Hospitals, including support to the iSEQ Center, the GenomeDK HPC facility, and the CIRRAU Center. Ethical approval for iPSYCH was provided by the Danish Scientific Ethics Committee, the Danish Health Data Authority, the Danish data protection agency and the Danish Neonatal Screening Biobank Steering Committee.

The research conducted using the GERA resource was approved as part of NIH project #21685 and data is available from dbGaP accession reference phs000674.v3.p3. We are grateful to GERA and the Kaiser Permanente Research Program on Genes, Environment, and Health (RPGEH) and its participants. Data came from a grant, the Resource for Genetic Epidemiology Research in Adult Health and Aging (RC2 AG033067; Schaefer and Risch, PIs) awarded to the RPGEH and the UCSF Institute for Human Genetics. The RPGEH was supported by grants from the Robert Wood Johnson Foundation, the Wayne and Gladys Valley Foundation, the Ellison Medical Foundation, Kaiser Permanente Northern California, and the Kaiser Permanente National and Northern California Community Benefit Programs.

DMH is supported by a Sir Henry Wellcome Postdoctoral Fellowship (Reference 213674/Z/18/Z) and a 2018 NARSAD Young Investigator Grant from the Brain & Behavior Research Foundation (Ref: 27404). A.M.McI acknowledges support from the Wellcome Trust (Wellcome Trust Strategic Award ‘STratifying Resilience and Depression Longitudinally’ (STRADL) Reference 104036/Z/14/Z and The Sackler Trust. N.R.W. acknowledges NMHRC grants 1078901 and 1087889. C.M.L acknowledges MRC grant MR/N015746/1. S.P.H. acknowledges MRC grant MR/S0151132. This investigation represents independent research part-funded by the National Institute for Health Research (NIHR) Biomedical Research Centre at South London and Maudsley NHS Foundation Trust and King’s College London. The views expressed are those of the authors and not necessarily those of the NHS, the NIHR or the Department of Health.

